# Photoacoustic pigment relocalization sensor

**DOI:** 10.1101/455022

**Authors:** Antonella Lauri, Dominik Soliman, Murad Omar, Anja Stelzl, Vasilis Ntziachristos, Gil G. Westmeyer

## Abstract

Photoacoustic (optoacoustic) imaging can extract molecular information with deeper tissue penetration than possible by fluorescence microscopy techniques. However, there is currently still a lack of robust genetically controlled contrast agents and molecular sensors that can dynamically detect biological analytes of interest with photoacoustics. In this biomimetic approach, we took inspiration from cuttlefish who can change their color by relocalizing pigment-filled organelles in so-called chromatophore cells under neurohumoral control. Analogously, we tested the use of melanophore cells from *Xenopus laevis*, containing compartments (melanosomes) filled with strongly absorbing melanin, as whole-cell sensors for optoacoustic imaging. Our results show that pigment relocalization in these cells, which is dependent on binding of a ligand of interest to a specific G protein-coupled receptor (GPCR), can be monitored *in vitro* and *in vivo* using photoacoustic mesoscopy. In addition to changes in the photoacoustic signal amplitudes, we could furthermore detect the melanosome aggregation process by a change in the frequency content of the photoacoustic signals. Using bioinspired engineering, we thus introduce a photoacoustic pigment relocalization sensor (PaPiReS) for molecular photoacoustic imaging of GPCR-mediated signaling molecules.

## INTRODUCTION

Photoacoustic (optoacoustic) imaging is based on the photoacoustic effect – the conversion of absorbed light energy into heat, causing the generation of acoustic waves that can be captured by ultrasound detectors^1^. Over recent years, the optoacoustic imaging technique has been increasingly used in biomedicine because of its advantageous combination of optical and ultrasound imaging, decoupling the underlying mechanisms of contrast and spatial resolution. On the one hand, optoacoustic imaging relies on absorption-based contrast, allowing for rich optical differentiation of photoabsorbing molecules such as exogenous contrast agents or endogenous tissue chromophores^2^. On the other hand, optoacoustic imaging achieves penetration depths of millimeters or even centimeters and high ultrasonic resolutions because of the weak acoustic scattering in tissue.

In general, the acoustic signals measured in photoacoustics contain a broad range of ultrasonic frequencies. The size of the imaged photoabsorbers governs the frequency content of the photoacoustic signal: smaller absorbers generate higher photoacoustic frequencies than larger absorbers. For example, a spherical absorber with 20 *µ*m diameter generates frequencies up to ~100 MHz, while a 10 *µ*m absorber produces frequencies as high as ~200 MHz^3^.

Furthermore, high frequencies experience stronger acoustic attenuation than lower frequencies^4^, leading to an inverse relationship of highest measurable frequency and thus achievable spatial resolution with imaging depth.

Due to this relationship, optoacoustic imaging is a highly scalable modality regarding spatial resolution and imaging depth. Recently, we have introduced raster-scan optoacoustic mesoscopy (RSOM) in epi-illumination mode, offering imaging depths of several millimeters and a lateral and axial resolution of 20-30 *µ*m and ~5 *µ*m, respectively^5^, thereby bridging the gap between macroscopic and microscopic optoacoustic imaging modalities.

Up to now, contrast generation for optoacoustic imaging has mostly relied on utilizing endogenous chromophores or externally administered photoabsorbers such as organic dyes or nanoparticles^2^. With respect to genetically encoded contrast agents for optoacoustic imaging, over-expression of (reversibly switchable) fluorescent proteins that absorb in the visible or infrared range have been explored, in which either cyclized amino acids or cofactors such as biliverdin act as the chromophore ^6–9^.

Alternatively, biosynthetic pigments such as melanin and violacein have been utilized as contrast agents, which are beneficial because of their generally higher photobleaching resistance, enzymatic amplification of the genetic signal and potentially higher local concentrations as compared to chromoproteins^10,11^. The advantages of melanin are that it is synthesized in a monoenzymatic reaction and that it yields strong and broad absorption over the visible range extending into the near-infrared^12^. Moreover, melanin exhibits a fluorescence quantum yield of less than 1% thereby maximizing heat generation by non-radiative decay, a prerequisite for photoacoustic signal generation^12^. For cell types that endogenously produce and sequester melanin such as melanophores, it represents an excellent contrast agent for optoacoustic imaging that has, for instance, enabled volumetric tracking of melanophore migration during zebrafish development^13^. If melanin production is not compartmentalized however, it has been shown to be toxic in standard cell lines^14^. In melanophores, melanin is stored in specialized organelles that are transported inside the cell along its microtubular cytoskeletal network, referred to as melanosomes.

Certain animals such as chameleons and cuttlefish are able to adjust their skin color to the environment by translocating the melanosomes within melanophores, which is mediated by neuronal and hormonal inputs^15^. This impressive behavior is used for camouflage to hide from predators and in the case of the cuttlefish may also be used as means of communication^16^. Forms of camouflage have been reported also in the common laboratory fish model zebrafish (*Danio rerio*). If a zebrafish is on a dark background, its melanophores disperse their melanosomes, leading to a darker appearance of the melanized fish skin. Conversely, aggregation of melanosomes around the nucleus of the melanophores triggered by bright background colors results in a lighter appearance of the animal^17^.

Classical neurohumoral molecules are involved in regulating this melanosome relocalization within melanophores. These include the pituitary-derived melanin-stimulating hormone (α-MSH), which induces melanosome dispersion, and the hypothalamus-derived melanin-concentrating hormone (MCH)^17-19^, which induces melanosome aggregation. Other hormones that influence melanosome translocation include melatonin^20,21^, catecholamines such as noradrenaline^15,22^, and serotonin^23^. The mechanism of action of these molecules is realized via binding to G-protein–coupled receptors (GPCR) expressed on the cell surface of melanophores, which modulates the intracellular levels of cyclic adenosine 3’-5’-monophosphate (cAMP) and calcium. These secondary messengers ultimately control melanosome relocalization inside the cell through actions on the cytoskeletal transport machinery^24,25^.

Inspired by the cellular melanosome aggregation and dispersion process and by the ideal photophysical properties of melanin, we were interested in how the reversible relocalization of strongly photoabsorbing pigments within melanophores could be exploited as a dynamic contrast mechanism for molecular optoacoustic imaging.

To this end, we selected an immortalized melanophore cell line from the frog *Xenopus laevis* that has been successfully used as a robust *in vitro* readout to study complex GPCR-mediated signaling transduction pathways^26^. Convenient plate-reader based monitoring of absorbance changes resulting from melanosome relocalization, this methodology contributed to the characterization of important neuroendocrine pathways, activated for example by the pineal hormone dopamine^26^ and melatonin^27^.

Both *Xenopus laevis* and zebrafish are cold-blooded animals and are cultured at similar temperatures. Therefore, we explored whether xenografting wildtype or genetically modified frog melanophore cell lines into zebrafish could be employed as a whole-cell photoacoustic sensor for GPCR ligands. In distinction to the well-known CNiFER sensors^28^ that detect a specific GPCR ligand such as glutamate via a downstream fluorescent calcium indicator signal in xenografted human embryonic kidney (HEK) cells, photoacoustics could directly image melanosome relocalization via changes in the photoacoustic signal amplitudes.

For gold and melanin-like nanoparticles, it has been demonstrated that an aggregation of the nanoparticles can lead to an increase of photoacoustic signal amplitude due to overlapping transient thermal fields surrounding the nanoparticles^29,30^. Changes of photoacoustic frequencies have also been measured in the context of detecting aggregation and morphological changes of red blood cells in microscopic settings *in vitro*^*31*,*32*^.

In the given context, we thus reasoned that dynamic pigment relocalization could also be detectable by shifts in the photoacoustic frequencies above 10 MHz because the average size of the imaged photoabsorbing objects is altered by the melanophore dynamics. A demonstration on intracellular pigment dynamics *in vivo* would establish the frequency content of photoacoustic signals as a new observable for the high-resolution photoacoustic sensing of dynamic molecular processes in live animals.

For the optoacoustic 3D monitoring of melanophore dynamics, we selected the RSOM modality as it combines a number of advantages over traditional optical imaging systems. First, RSOM achieves the necessary spatial resolution to resolve single melanophores and to track changes in the absorptive volume of the imaged cells. Second, RSOM relies on widefield laser illumination and acoustic focusing (as opposed to optical focusing), being less prone to pigment shadowing than pure optical modalities. Third, the photoacoustic frequencies detectable via RSOM (> 200 MHz) are high enough to capture frequency shifts generated by changes in absorber size in the range of approximately 10-40 *µ*m brought about by melanosome relocalization^17^.

Combining engineered melanophore cells with RSOM, we introduce here a new reversible contrast mechanism for molecular imaging with photoacoustics based on the relocalization of pigments in response to analytes of interest. Such a photoacoustic pigment relocalization sensor (PaPiReS) is distinct from sensors that change their color in response to analytes of interest^33-35^ because the absorbance spectrum of the pigments does not change. In addition, we could demonstrate how GPCR signaling can be mapped with optoacoustic imaging and how a change in the frequency content of photoacoustic signals can be used as a reversible sensor readout.

## RESULTS

### Melanophore aggregation during background adaptation and upon MCH treatment in zebrafish larvae can be detected with RSOM

To test the feasibility of detecting GPCR-mediated pigment relocalization in engineered melanophores, we first asked whether optoacoustic mesoscopy could detect naturally occurring pigment relocalization in melanophores during background adaptation of zebrafish. To this end, we induced pigment dispersion in melanophores of zebrafish larvae by adapting them to a dark environment for ~16 hours. Subsequently, we kept the larvae on a black background while imaging them using epi-illumination RSOM (Figure 1a; Supplementary Figure S1). Exposing the fish to a combination of a white background and light illumination for 30 minutes triggered melanophore aggregation as observed with RSOM (Figure 1b). Because of their high content of melanin, eyes were easily recognized in the photoacoustic images, which were obtained by projecting the 3D reconstructions of the fish along the depth dimension (*z*-axis), representing maximum intensity projections (MIP) in top view (Figure 1a,b). As expected, melanophores along the body also produced good photoacoustic contrast. Several stripes were visible in the photoacoustic image of the fish in the control state comprising identifiable melanophore clusters or single cells (Figure 1a, white arrows in the inset).

**Figure 1.**
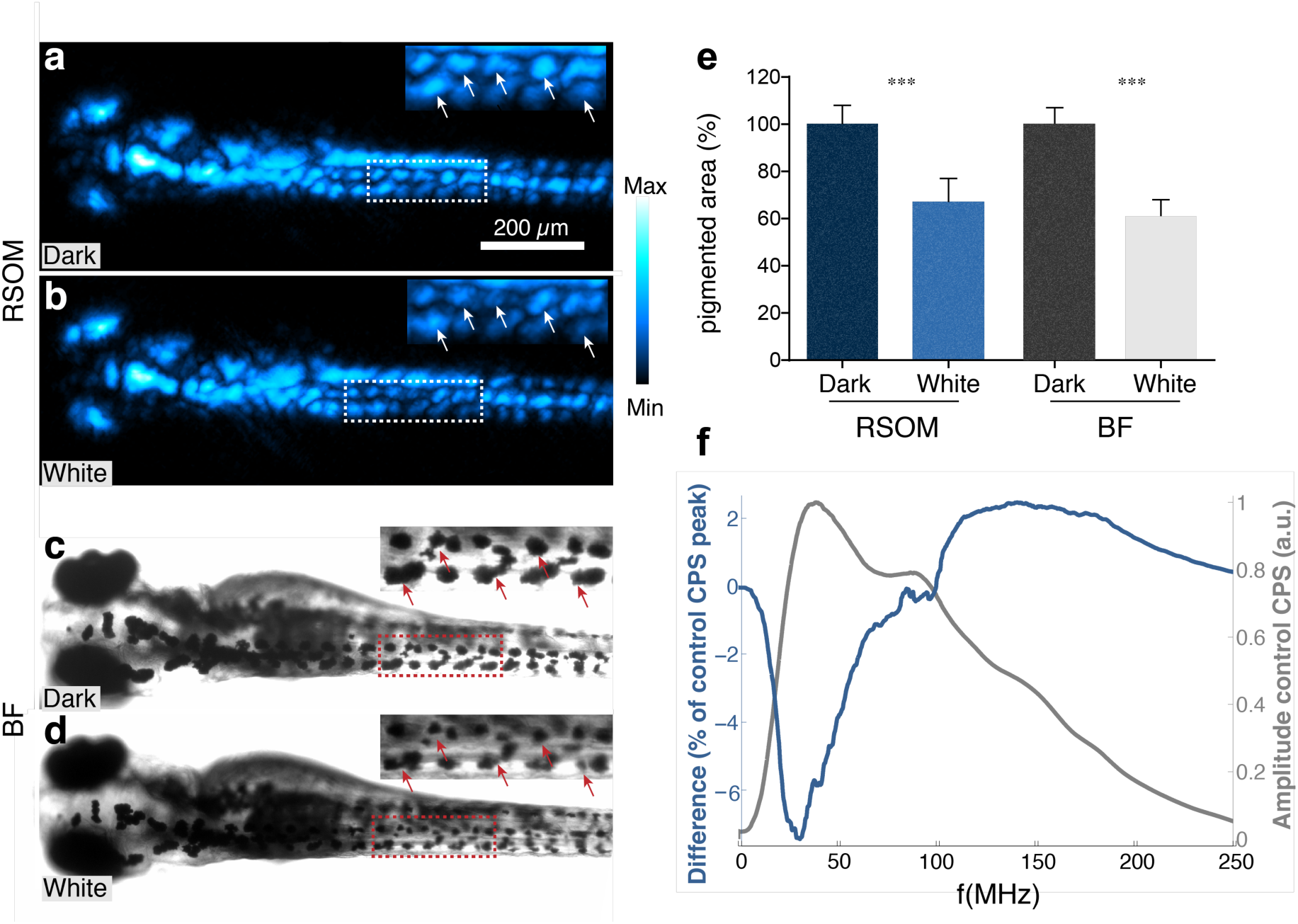
RSOM imaging and photoacoustic signal frequency analysis of melanophores after background adaptation of a zebrafish larva *in vivo.* **(a,b)** Top-view maximum intensity projections (MIP) of a 5-day-old zebrafish larva imaged with RSOM after being adapted to a dark **(a)** or a light **(b)** environment. Dorsal view, anterior is left. The insets show a magnification of the trunk region delineated by the white dashed boxes. White arrows highlight positions in which background-dependent melanosome aggregation is observed. **(c,d)** Corresponding brightfield (BF) microscopy images of the same fish as shown in (a) and (b) obtained after optoacoustic imaging. The insets show a magnification of the regions indicated by the red dashed outline, corresponding to the insets in (a) and (b). Red arrows mark the same cells containing aggregated melanosomes as labeled in (a) and (b). **(e)** Quantification of the melanosome aggregation effect for the two conditions and imaging modalities shown in (a)-(d) (mean pigmented area normalized to the maximum of each modality ± SEM from 19 cells (BF) or 9 clusters of cells (RSOM)). A significantly reduced pigmented area was observed after light background adaptation with both RSOM and BF (paired t-test, *** indicates p < 0.001). **(f)** Difference in photoacoustic signal cumulative power spectra (CPS) normalized to the CPS peak value of the control measurement. The light blue curve shows the difference between the CPS obtained from the light background adapted fish and the dark-adapted fish. The gray curve represents the CPS obtained from the dark-adapted fish.

In fish adapted to a white background, regions with a prominent reduction in the pigmented area could be distinguished in the photoacoustic images (white arrows in the insets of Figure 1b as compared to Figure 1a). Moreover, because of the deep tissue penetration achieved by RSOM, we were able to observe melanosome aggregation also at different depths and from different angles of the three-dimensional reconstruction. This allowed us to discern pigment patterns otherwise disguised in the single depth-projection images. A full 360° rotation of the 3D reconstruction of the imaged zebrafish before stimulation is shown in Supplementary Movie S1. Subsequent to the photoacoustic measurement session, we repeated the background adaptation experiments with the same fish under a brightfield (BF) microscope (Figure 1c,d). Even though the low contrast of the brightfield images prevented a detailed study of the melanosome aggregation effect, we could identify responding melanophores at similar locations along the anterior-posterior (A-P) axis of the fish as in the photoacoustic images (insets in Figure 1c,d).

To quantify and compare the melanophore aggregation response as captured by photoacoustic and brightfield imaging, we performed a manual segmentation of the regions shown in the respective insets (Figure 1a-d), which yielded a significant reduction in the mean pigmented area by 38% (± 3.1%, standard error of the mean (SEM)) due to white background adaptation under the brightfield microscope (paired t-test, *p*-value: 7.6× 10^-8^, *n*=19) and by 33% (± 4.7%) under RSOM imaging (paired t-test, *p*-value: 0.0003, *n*=9; Figure 1e). These data demonstrate that the imaging conditions in RSOM are sufficiently gentle and do not interfere with the background adaptation behavior of the fish.

Generally, smaller photoabsorbers should generate higher photoacoustic frequencies. Therefore, we set out to assess next, whether the dynamic process of melanosome aggregation is reflected in the frequency spectra of the recorded photoacoustic data. First, to study the effect of the melanosome aggregation state on photoacoustic frequencies, we carried out simulations of contracting and expanding absorbers and calculated the difference spectra of the resulting photoacoustic frequency distributions (Supplementary Figure S4). In the case of contracting absorbers, the difference spectra yielded distinct bipolar shapes with a low-frequency minimum and a high-frequency peak (Supplementary Figure S4a), whereas the shapes corresponding to expanding absorbers are inverted (Supplementary Figure S4b). Hence, melanosome aggregation is expected to be reflected by a distinct bipolar difference spectrum of photoacoustic frequencies.

In the next step, we performed a frequency analysis of the photoacoustic data recorded by RSOM from the zebrafish during background adaptation. To this end, we summed up the frequency spectra of all potentially melanin-related signals across the entire photoacoustic dataset to obtain the cumulative power spectra (CPS) for both light-adapted or dark-adapted conditions. The CPS of the fish measured in darkness exhibited a maximum at ~38 MHz (grey curve in Figure 1f). The difference spectrum of the CPS relative to the white and the dark background adapted fish showed a negative peak at ~30 MHz and a high-frequency excess above ~100 MHz (represented by the light blue curve in Figure 1f). The shape of the CPS difference spectrum obtained after the background adaptation experiment indicates a shift from low to high frequencies, as a result of a reduction in the average absorber size present in the living specimens.

Whereas these changes in the global frequency distributions were obtained from the entire scanned volume, we were next interested whether different regions along the A-P axis with strong frequency shifts corresponded to regions with a prominent reduction of the pigmented areas. Therefore, we mapped out the magnitude of melanophore aggregation via a position-dependent analysis of the amplitude and frequency-shift information of the photoacoustic signals. To this end, we segmented melanophore clusters in the trunk region in the dark and white background adapted states of the imaged zebrafish larva and compared the pigmented areas. For visualization, we colored the segmented clusters of the dark-adapted fish according to the magnitude of area change such that red, orange and yellow colors correspond to a change of more than 10%, 23%, and 30%, respectively (Supplementary Figure S2a). To visualize position-dependent frequency shift magnitudes, we performed the above-described frequency analysis at each position in the *xy*-plane (Supplementary Figure S2b,c) of the reconstructions for a neighborhood of 13 × 13 *z*-signals. The peak values of the CPS difference spectra are displayed in the frequency shift map in Supplementary Figure S2b (as compared to the photoacoustic MIP of the dark-adapted larva in Figure 1).

We noted that the most significant contribution of frequency shifts originated from the trunk region, whereas only little changes occurred in the head region. Furthermore, we could identify three regions of strong frequency change, as well as a larger area of weaker frequency shift (white and red arrows in Supplementary Figure S2b, respectively). Because single cells or small cell clusters were not easily detectable in the head region, segmentation was not performed there (*‘not segm.’* in Supplementary Figure S2a). When comparing the maps in the trunk region, a correlation between higher and lower magnitudes of frequency shift and stronger and weaker area changes could be identified. The clusters showing a reduction in the pigmented area of more than 23% and 30% were the ones that likely contributed the most to the global frequency shift observed after the adaptation to a white background (Figure 1).

The experiments carried out in zebrafish larvae showed that (i) RSOM can detect the dynamic process of melanin relocalization naturally occurring inside melanophores of live larvae (Figure 1a,b, Figure 2c,d, Supplementary Figure 2), and that (ii) RSOM acquisition does not interfere with the reversible pigment clustering process as shown by brightfield imaging of the same individual after the RSOM measurement (Figure 1c,d).

**Figure 2.**
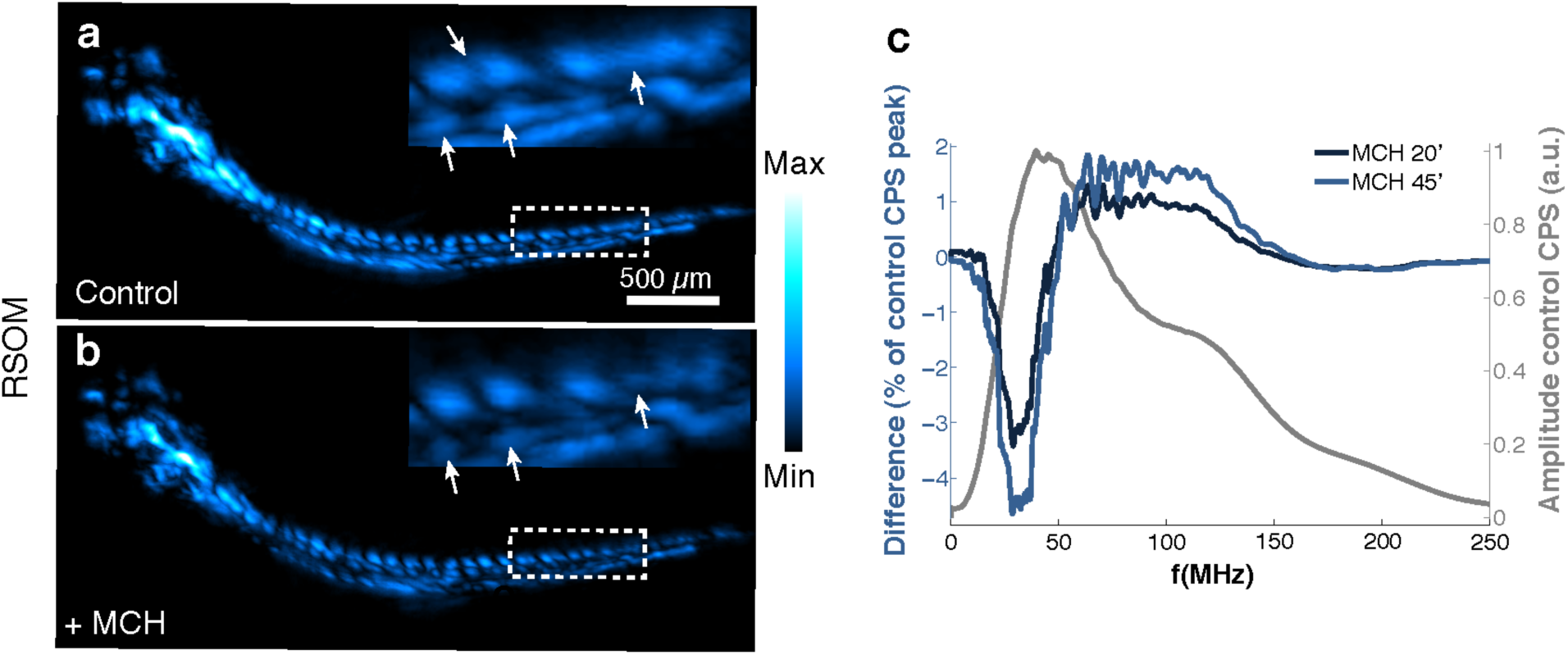
RSOM imaging of a zebrafish larva before and after exposure to MCH. **(a,b)** Top-view MIPs of a zebrafish larva imaged with RSOM before **(a)** and after **(b)** 45 minutes of exposure to 100 *µ*M of MCH dissolved in fish water. Dorsal view, anterior is left. The insets show a magnification of the trunk region delineated by the white dashed boxes. White arrows highlight regions of prominent melanosome aggregation. **(c)** Differences in photoacoustic cumulative power spectra (CPS) normalized to the control CPS peak value. The dark blue curve shows the difference between the CPS obtained after 20 minutes of treatment with MCH and the CPS of the control condition. The light blue curve shows the same analysis for 45 minutes of treatment with MCH. The CPS of the control condition is shown in gray color as a reference.

Next, to investigate whether hormone-induced melanophore aggregation could also be detected using RSOM, we tested larvae pigmentation upon delivery of MCH on live zebrafish, which is known to induce a visible aggregation of melanosomes in these animals^36^. Upon stimulation with MCH, the imaged melanophore size was reduced by 19% (± 4.2%) (white arrows in the insets of Figure 2a,b, paired *t*-test, *p*-value <0.001, *n*=13).

Again, we tested whether a global frequency change was discernible before and after either 20 or 45 minutes of exposure to MCH. We observed that the CPS of the control condition featured a maximum at ~45 MHz (grey curve in Figure 2c). The difference spectra of the CPS for the two different durations of MCH treatment (control condition subtracted) are illustrated in Figure 2c by the dark blue curve (20 minutes of MCH treatment) and the light blue curve (45 minutes of MCH treatment). Interestingly, both curves show a pronounced negative peak at ~30 MHz and a positive plateau in the 50-140 MHz range, indicating a shift from low to high frequencies upon melanosome aggregation. Furthermore, we noted that the longer MCH treatment curve exhibits a similar frequency profile, albeit with a higher peak-to-peak amplitude, indicating a stronger aggregation response due to the longer stimulation time.

### RSOM imaging of hormone-triggered melanosome aggregation in cultured melanophores

Next, we set out to investigate whether GPCR-mediated aggregation of melanosomes and the subsequent reduction of the pigmented area observable via optoacoustic mesoscopy could be employed to resolve melanophore responses in deep tissue. This capability could then be used to image xenotransplanted cells that could be genetically engineered to function as a whole-cell sensor for photoacoustics, similarly to the catecholamine sensors for light microscopy developed by the Kleinfeld laboratory^28^. Hence, we first explored the RSOM detection of melanosome aggregation in a well-established immortalized cell line from *Xenopus laevis.* These cells exhibit relatively large sizes in the range of 50-100 *µ*m and have been previously employed as a cell-based sensor using GPCR signaling *in vitro* by simple monitoring of absorbance changes in a plate reader format^26,37^. Exposure to MCH induced a significant melanosome aggregation response of frog melanophores, as seen in brightfield microscopy (Figure 3a-d), with the majority of melanosomes relocating around the nucleus. A low density of the frog melanophores was used such that individual cells could be differentiated for convenient segmentation and quantification of the reduction in the mean pigmented area, which was found to be 58% (± 5.3%) (Wilcoxon matched-paired signed rank test, *p*-value: 2×10^-5^, *n*=25).

**Figure 3.**
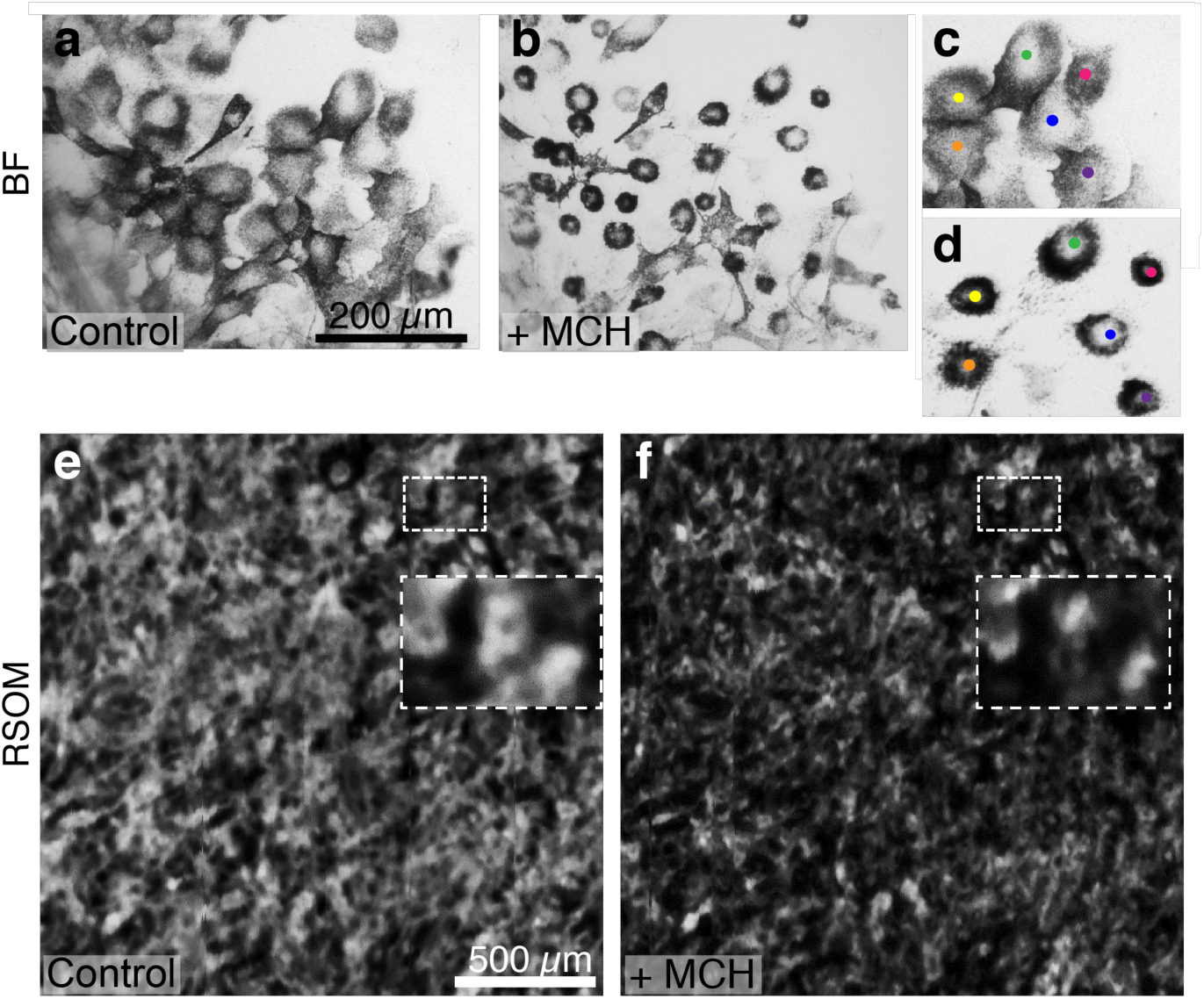
Cultured frog melanophores imaged with brightfield microscopy and RSOM before and after exposure to MCH. **(a-d)** Brightfield microscopy images showing cultured frog melanophores before **(a)** and after **(b)** 15 minutes of treatment with 30 *µ*M of MCH. **(c,d)** Close-ups of a subset of cells shown in (a) and (b). The cell identities are indicated with colored dots. A reduced pigmented area by 58% is observed as compared to the control image (Wilcoxon matched-paired signed rank test, p-value < 0.0001, n=25). **(e,f)** RSOM imaging of cultured frog melanophores before **(e)** and after **(f)** exposure to 30 *µ*M of MCH, which triggered melanosome aggregation. The images show top-view MIPs of the RSOM measurement of the cell layer, while the insets represent magnifications of the regions delineated by the dashed boxes. Upon MCH stimulation, the overall pigmented area is found to be reduced by 22% as compared to the control measurement (paired t-test, p-value: 0.006, n=10).

Using the same parameters, we next imaged the frog melanophores with RSOM and could clearly differentiate the dispersed state (vehicle control, Figure 3e) from the aggregated state after MCH stimulation (Figure 3f; reduction in size by 22% (± 5.5%), paired t-test, *p*-value: 0.006, *n*=10; please note that the contrast is inverted as compared to the BF images because highly absorbing structures give strong photoacoustic signals.) Remarkably, the resolution of the RSOM system was sufficient to even distinguish single frog melanophores and their nuclei as seen by the absence of signal in the center of the cells (insets in Figure 3e,f).

### Wildtype frog melanophores xenotransplanted into the brains of zebrafish respond to melatonin

Encouraged by the clear contrast changes observed by RSOM *in vitro*, we next sought to test whether GPCR-mediated pigment relocalization in frog melanophores could also be detected *in vivo*. We thus transplanted cultured frog cells into the midbrain of juvenile zebrafish. To ensure engraftment of the cells, we performed the transplantation in immunocompromised fish mutants (homozygous *rag2^E450fs^* on a *Casper* background^38^).

As can be seen in the photoacoustic top-view MIP of the imaged fish (Figure 4a), the individual xenotransplanted cells (orange color map) could clearly be differentiated from each other and from endogenous structures of the fish such as the pigmented eyes (blue color map) virtue of the absence of photoacoustic background signals in the head region. The identification of transplanted cells and endogenous structures (*e.g.,* fish eyes) in the photoacoustic images was also confirmed by the corresponding BF image of the head region of the same fish (inset in Figure 4a). After stimulating the xenotransplanted melanophores for 30 minutes with melatonin, we were able to pick up a melanosome aggregation response of the cells by observing a clear reduction of their pigmented areas by 30% (± 9.5%) in the photoacoustic images (Figure 4b-c, paired t-test, *p*-value: 0.008, *n*=12).

**Figure 4.**
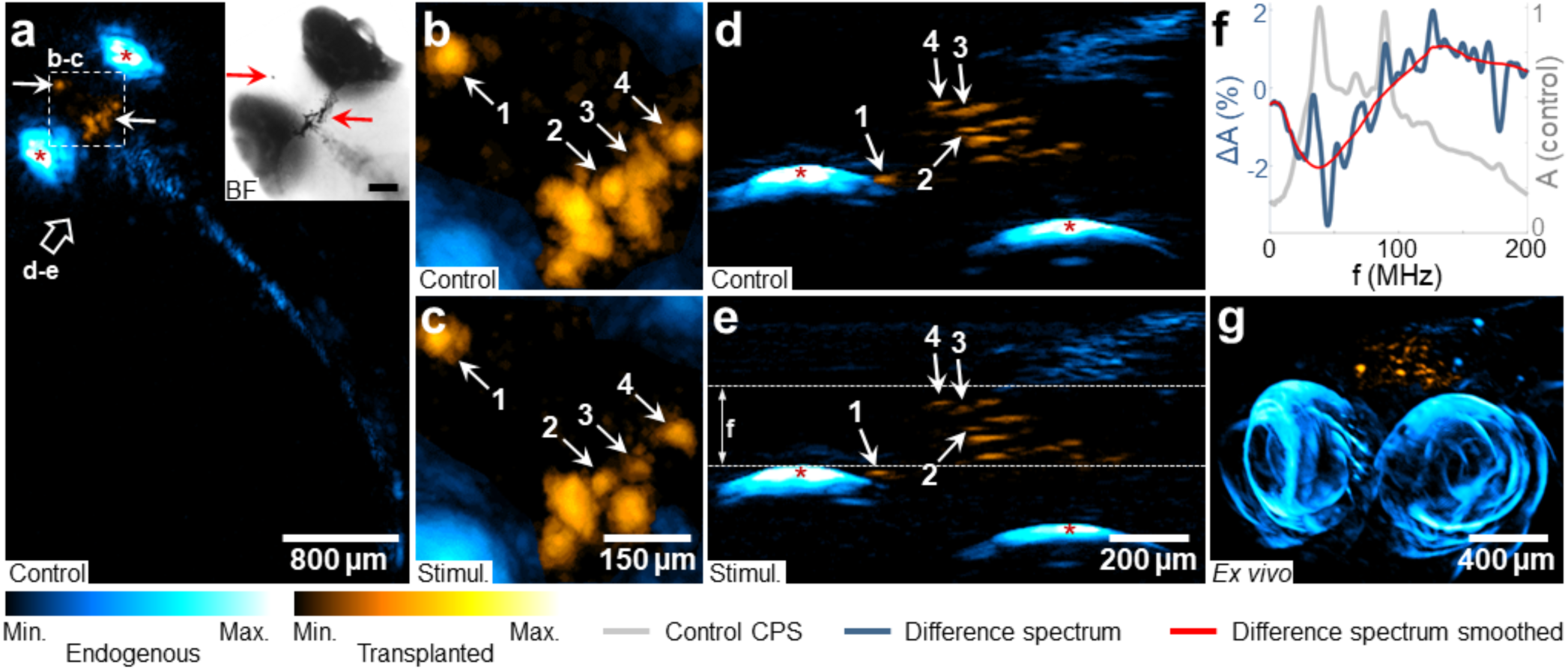
Optoacoustic imaging of melatonin-induced melanosome aggregation in xenotransplanted frog melanophores in zebrafish *in vivo*. **(a)** Xenografted frog melanophores (orange color map) in *rag2^E450fs^* mutant fish. Shown is a top-view MIP of the RSOM measurement of the fish before stimulation (‘Control’). Background signals from endogenous pigmentation are displayed with a blue color map, whereas the eyes are additionally marked with red asterisks. The inset shows a corresponding brightfield image of the head region, which was recorded before RSOM imaging. Scale bar: 200 *µ*m. **(b,c)** Zoom into the region indicated by the white dashed box in (a). Four melanophores are labeled for reference and shown before (b) and after (c) 30 min of stimulation with 0.1 *µ*M of melatonin (‘Stimul.’). **(d,e)** Side-view MIPs of the RSOM reconstructions in a direction indicated by the white arrow in (a). The gray dashed lines in (e) indicate the part of the reconstructed volumes that was subjected to the photoacoustic frequency analysis. **(f)** Difference of cumulative power spectra (CPS) (Stimul. minus control; blue curve) normalized to the maximum of the control CPS (gray curve). The red curve represents a local regression fit to the blue spectrum to highlight the global trend. **(g)** MORSOM imaging of the same specimen *ex vivo* after the stimulation experiment showing a side-view MIP of the 3D reconstruction. Cyan color: native cells and eyes. Orange color: region around the xenotransplanted frog cells.

Furthermore, the tomographic capabilities of the RSOM technique enabled us to study the triggered pigment relocalization effect in 3D. The cells appeared distributed throughout a depth of ~300 *µ*m of the fish brain (90° side views, Figure 4d-e). Without the melatonin stimulation, no change in the imaged cell volumes was observed in repeated control images of a single transplanted *Xenopus* melanophore in a living zebrafish larva. Furthermore, consecutive measurements of a single transplanted 10 *µ*m black polystyrene microsphere did not show any substantial changes in the images, demonstrating the stability of the imaging system (Supplementary Figure S3).

In addition to analyzing the signal amplitudes in the photoacoustic images, we performed a frequency analysis of the photoacoustic signals derived from the xenotransplanted melanophores. The difference spectrum obtained from the CPS of the volume delineated by the white dashed lines in Figure 4e during the control and melatonin-stimulated condition (control spectrum subtracted) is shown as a blue curve in Figure 4f. A local regression fit is plotted in red color, while the CPS of the measurement before stimulation is shown in gray color as a reference. The difference spectrum exhibits a clear bipolar shape with a low-frequency minimum and a high-frequency excess, indicating a shift towards higher photoacoustic frequencies as a result of the stimulated melanosome aggregation. Subsequent to the RSOM imaging, the fish was scanned using a multi-orientation implementation of the RSOM technique (MORSOM)^39^, serving as a high-fidelity anatomical reference (Figure 4g). In MORSOM, the sample is rotated and scanned with the RSOM technique repeatedly from multiple angles, yielding an isotropic spatial resolution orthogonal to the sample rotation axis. This resulted in an improved 3D visualization of the imaged structures (a 360° rotation of the reconstructed specimen imaged with MORSOM is shown in Supplementary Movie S2).

### Engineered whole-cell photoacoustic sensor for an orthogonal GPCR ligand

After having established that RSOM achieves sufficient spatial resolution and sensitivity using xenotransplanted melanophores responding to sub-micromolar concentrations of melatonin in zebrafish, we were next interested whether this contrast mechanism could be generalized to other GPCR-ligands. We, therefore, overexpressed the designer receptor hMD4i-2A-eGFP (designer receptor exclusively activated by designer drugs, (DREADD) ^40,41^ in frog melanophores. hMD4i is a Gi-coupled receptor, engineered such that, when activated by the orthogonal agonist clozapine N-oxide (CNO), it causes a reduction of the intracellular levels of cAMP^40^. Given the physiological response of the melanophores to the Gi signaling^24,25^, we predicted that activation of this receptor should induce pigment aggregation in our cultured melanophores (Figure 5a), as was indeed observed *in vitro* (Figure 5b). In a proof of principle experiment, cells expressing the hMD4i-2A-eGFP were selected and enriched via fluorescence-activated cell sorting (FACS) (‘p4’ in Figure 5c) and subsequently xenotransplanted into *rag2^E450fs^* mutant zebrafish. We could again detect the engineered engrafted hMD4i-2A-eGFP+ cells in the head region (Figure 5d,e, orange color map, green dotted boxes) with very low background signals in RSOM. Subsequent administration of CNO then resulted in a reduction of the melanized area of the transplanted cells by 13% (± 5.6%) mirroring the response observed *in vitro,* although statistical significance was not reached (paired t-test, *p*-value: 0.39, *n*=3) due to the small number of cells implanted.

**Figure 5.**
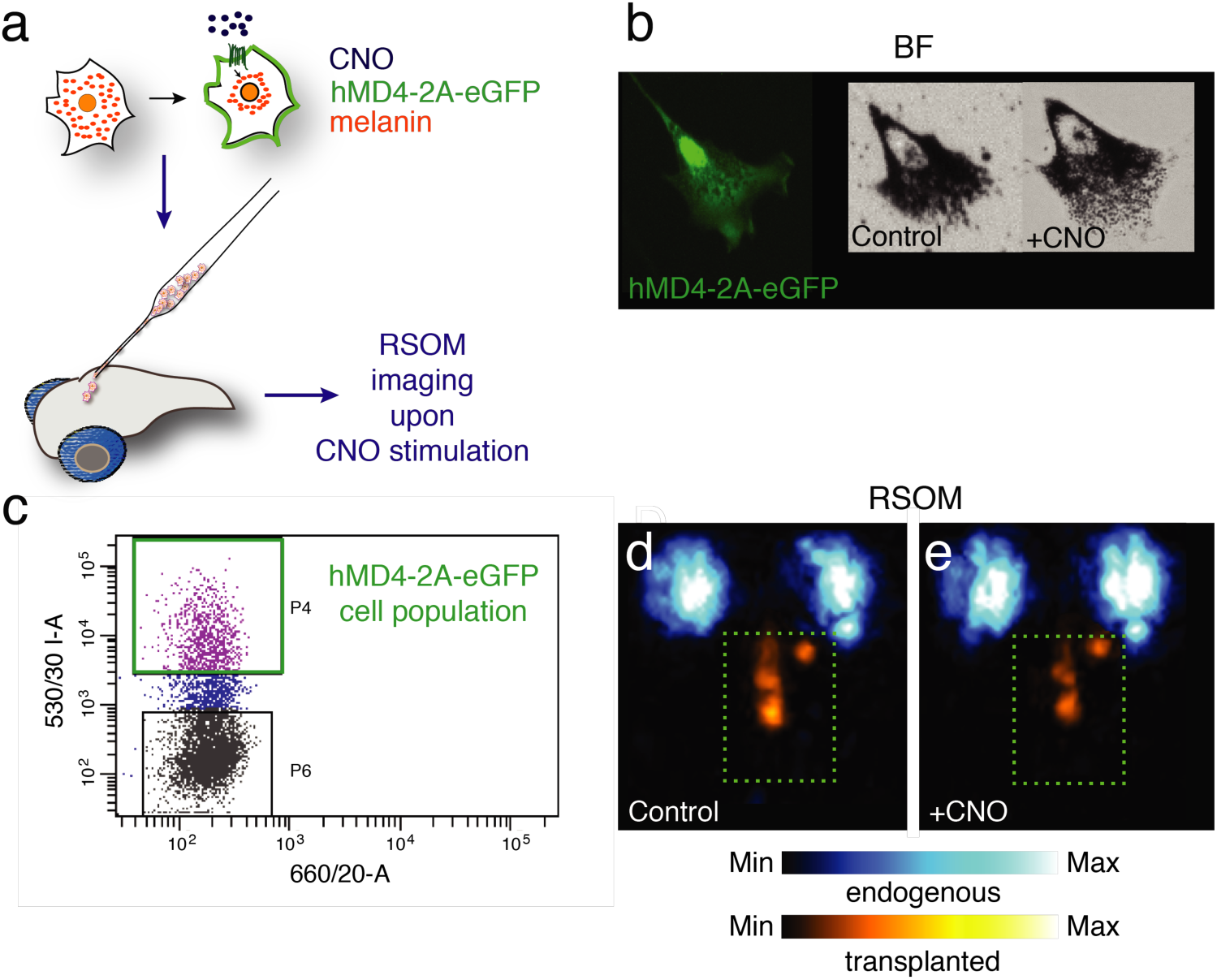
RSOM imaging of *Xenopus* melanophores expressing a designer receptor exclusively activated by designer drugs (DREADD) hMD4i xenografted into *rag2^E450fs^* mutant zebrafish. **(a-c)** Schematic showing the experimental design comprising co-expression of the DREADD hMD4i and the enhanced GFP (hMD4i-2A-eGFP) to enable fluorescence-activated cell sorting (FACS) of the most strongly expressing cells. **(b)** Brightfield imaging of melanosome aggregation in response to clozapine-N-oxide (CNO) stimulation of the transplanted melanophores expressing the hMD4i-2A-eGFP. (c) Dot blot showing the cell population selected by FACS cell sorting via eGFP fluorescence. **(d,e)** Photoacoustic top-view MIPs of a *rag2^E450fs^* mutant zebrafish with xenografted genetically modified frog melanophores (hMD4-2A-eGFP, orange color map) before **(d)** and after **(e)** stimulation with CNO. Cyan color map: endogenous structures and fish eyes. Orange color map: region around the xenotransplanted frog melanophores.

## DISCUSSION

We have shown that melanophore dynamics can be mapped *in vivo* by optoacoustic mesoscopy non-invasively in deep tissue with cellular resolution. In addition to the high contrast derived from the photoacoustic signal amplitudes, we could also demonstrate that the frequency content of the photoacoustic signals can be utilized to identify the activation states of natural and engineered melanophores *in vivo*.

The epi-illumination RSOM modality employed in this work has been proven to be particularly suitable for the non-invasive monitoring of 3D melanophore aggregation in living zebrafish. RSOM yielded sufficient contrast and spatial resolution to visualize individual melanophores in 3D and was confirmed to not impede melanophore activity of zebrafish during imaging. The analysis of photoacoustic frequency shifts consistently revealed the occurrence of melanosome aggregation in the imaged specimens, both in measurements of the entire imaging volume and in a spatially resolved manner. The frequency of the photoacoustic signal could, therefore, constitute a valuable additional observable for studying dynamic cellular processes via photoacoustics *in vivo*. We have furthermore complemented the RSOM imaging with the recently developed MORSOM approach that yielded isotropic spatial resolution in the plane perpendicular to the sample rotation axis, thus facilitating the anatomical identification of transplanted cells albeit at the cost of a longer acquisition time.

After establishing the suitability of RSOM for monitoring 3D melanophore dynamics in living fish, we used it to demonstrate a new biomimetic sensor mechanism based on dynamic pigment relocalization within bioengineered melanophores. The dynamic cell sensors provided robust photoacoustic contrast due to the strong absorbance and high photobleaching resistance of melanin. We showcased this mechanism by the detection of melatonin, which is a critical molecular signaling molecule involved in circadian rhythm. We furthermore provided a proof of principle on how this sensor mechanism could be generalized to other GPCR ligands by genetically installing a response to the orthogonal DREADD ligand clozapine. Responses to other GPCR ligands of interest could be engineered analogously by expressing the respective GPCRs and ablating undesired GPCR inputs. Instead of xenografting melanophores into host organisms, melanophores could in principle also be genetically modified *in situ* to be responsive in a host organism such as zebrafish. It is also conceivable that the pigment relocalization mechanism is generalized to other contractile cells, such as smooth muscle cells. The principle could alternatively be realized in a system consisting of synthetic or genetically encoded photoabsorbing nanostructures functionalized on their surface with moieties that change their agglomeration state as a function of an analyte of interest.

In summary, we introduced a photoacoustic relocalization sensor (PaPiReS) for molecular photoacoustic imaging based on the dynamic relocalization of pigments with a fixed absorption spectrum as opposed to a complementary mechanism based on a color change of photoabsorbing molecules. We could also demonstrate how the frequency content in photoacoustic signals can be utilized to derive spatially resolved information about melanophore dynamics *in vivo* to obtain molecular information with optoacoustic imaging. This work furthermore constitutes the first demonstration of a photoacoustic sensor for signal transduction via the important class of GPCRs. The biomimetic sensor mechanism disclosed herein can be generalized to other ligands or stimuli and different cell systems with mobile chromophores. PaPiReS could be used to multiplex via spectral decomposition of different pigments and could also be combined with sensors based on chromic agents to substantially enrich the palette of molecular sensors for optoacoustic imaging.

## MATERIALS AND METHODS

### Frog melanophores cell culture and hormonal treatments

Immortalized melanophores from *Xenopus Laevis* (B27 cell line, a kind gift from Dr. M. Lerner, Harvard Medical School) were propagated as previously described^37^ in fibroblast-conditioned medium (CFM) containing 50 *µ*g/ml of insulin (Sigma Aldrich). For the imaging experiments, the cells were seeded in 35 mm tissue culture plates and cultured for 8-10 days before the experiment. The cells were washed in 0.7x phosphate buffer saline (PBS, Life Technologies, Carlsbad, CA, US) and then starved in 0.7x L15 medium (Sigma Aldrich, St. Louis, MO, USA) between 10-14 hours prior to imaging. To induce melanosome aggregation, cells were washed twice in PBS and subsequently treated with 30 *µ*M of rat MCH for 40 minutes.

### Zebrafish maintenance and preparation

Zebrafish larvae were bred and maintained according to standard conditions^42^. The fertilized eggs from wildtype AB strain *Casper* or *rag2^E450fs^* mutant fish were raised in embryo medium at 28°C. All procedures involving animals and their care conformed to the institutional guidelines.

### Background adaptation experiment in zebrafish larvae

5-day-old larvae were raised in the dark for ~16 hours and kept in the dark during the mounting procedure and prior to RSOM imaging. For imaging, the fish were embedded in 1% low melting agarose. Two fish were scanned simultaneously. First, the specimens were imaged in darkness and in front of a black background. Subsequently, the fish were exposed to bright light for 30 minutes while facing a white background. This condition was kept during the second RSOM scan.

### MCH mediated melanosome aggregation experiments in zebrafish larvae

The larvae were raised in a standard light/dark cycle and embedded in 1% low melting agarose. To elicit melanophore aggregation, the fish were treated with 100 *µ*M of rat MCH (Sigma Aldrich) and imaged after 20 and 45 minutes of hormone incubation. As a control, untreated fish were imaged two consecutive times.

### HmD4i-2A-eGFP plasmid preparation and DNA transduction of frog melanophores

The hmD4i cassettes was excised by standard PCR from the vector pAAV-hSyn-DIO-HA-hM4D(Gi)-IRES-mCitrine (Addgene Plasmid #50455). The cassette was cloned in frame with a 2A-eGFP (obtained by standard PCR from a donor vector) using Gibson Assembly (NEB, New England Biolabs, Ipswich, MA, USA). *Xenopus* melanophores were freshly seeded two days before DNA transduction. 4 *µ*g of purified plasmid were transduced into frog melanophore cells using electroporation with *‘* Ingenio Electroporation Solution’ (Mirus Bio LLC, Madison, WI, USA) and *‘* Gene Pulser Xcell™’ (Biorad Laboratories, Hercules, CA, USA) with the following settings: 250 V, 800 *µ*F, 700 Ohms. The cells were left to recover and their gene expression was assessed via fluorescence at 48 hours after the electroporation. Cells were selected via FACSAria III (BD Bioscience, Franklin Lakes, NJ, USA) 3 days after the electroporation and the results were analyzed with ‘Flowing Software’.

### Xenotransplantation of frog melanophores and beads in zebrafish

Frog melanophore cells were collected in PBS via centrifugation and were xenotransplanted - using standard pressure injection- into the brain of a *rag2^E450fs^* mutant fish, which were anesthetized in MS222 (PharmaQ, Overhalla, Norway) treated with a local analgesic. After surgery, the fish were left to recover for at least 16 hours. Successful engraftment of the transplanted cells was observed under a standard stereomicroscope revealing how the melanophores spreaded their processes into the brain tissue. The fish were then imaged with RSOM and euthanized subsequently using a high dose of MS222.

### Melatonin-mediated melanosome aggregation experiment

One day after xenotransplantation of the frog melanophores, the fish were imaged with RSOM before and after addition of 100 nM of melatonin (Sigma Aldrich) to the bath. The fish were scanned before and 30 minutes after the hormone was delivered, similarly to the MCH experiment.

### Optoacoustic imaging procedure

Optoacoustic imaging was performed by means of epi-illumination raster-scan optoacoustic mesoscopy (RSOM)^5^. A scheme of the imaging system is shown in Supplementary Figure S1. The samples were illuminated by a pulsed solid-state laser at a wavelength of 532 nm (Wedge HB532, Brightsolutions SRL, Pavia, Italy; actively Q-switched, pulse width: ~1 ns, max. pulse repetition rate: 2 kHz, max. energy per pulse: 1 mJ.) The laser beam was expanded and coupled into an optical fiber bundle with three arms (1808B, Ceram Optec GmbH, Bonn, Germany; NA: 0.22) to guide the illumination onto the sample from the same side as the detection was performed (epiillumination). The acoustic waves generated in the sample were detected by a spherically focused ultrasound transducer at a central frequency of ~100 MHz (SONAXIS, Besancon, France; -12 dB bandwidth (pulse-echo method): ~20-180 MHz, focal distance: 1.6 mm, active element diameter: 1.5 mm). By measuring a photoacoustic point source (receive-only method), a -18 dB high cutoff frequency of ~225 MHz of the transducer has been determined experimentally. Due to the short focal distance of the transducer, the front end was conically shaped to allow the illumination to be brought underneath the transducer at an angle of approximately 45° (see Supplementary Figure S1). In order to provide efficient acoustic coupling, the transducer was immersed in the liquid covering the sample during the respective measurement (*e.g.,* fish water, PBS or MCH). The focal region of the transducer was positioned above the samples or structures of interest to be imaged, *i.e.,* the photoacoustic signals originated from the conical far-field of the spherically focused detector. After amplification with a low-noise amplifier (AU-1291, Miteq Inc, USA; gain: 63 dB), the detected photoacoustic signals were recorded by a high-speed digitizer at a sampling rate of 1GS/s (EON-121-G20, Gage-applied, Montreal, Canada; vertical resolution: 12 bit). For the light adaptation experiment, the signals were acquired with a similar device at 0.9 GS/s (ADQ-412-3G, SP Devices, Linköping, Sweden; vertical resolution: 12 bit). The trigger for the data acquisition was provided by a photodiode (DET36A, Thorlabs, Newton, NJ, USA) that collected scattered laser light. Raster-scanning was performed by laterally moving the transducer together with the illumination over the sample in a semi-continuous manner by means of two motorized high-precision piezo stages (M-683.2U4, Physik Instrumente GmbH & Co. KG, Karlsruhe, Germany). Along the continuous fast axis, the step size between consecutive measurements was determined by the laser repetition rate and the speed of the fast axis. The step size was 10 *µ*m for the melatonin stimulation experiment yielding an acquisition time of fewer than 4 minutes for a scanning field of view (FOV) of 6 × 6 mm^2^. For the other experiments, the step size was reduced to 4 *µ*m to improve the signal-to-noise ratio (SNR), resulting in an acquisition time of ~10 minutes for a 6 × 6 mm^2^ FOV. At each measurement position, the time-resolved photoacoustic signals were recorded, bandpass filtered in the 20-180 MHz range to reject noise and stored for further processing. Parasitic signals originating from laser light reflections were removed using a spatial FFT filter.

### Image reconstruction

RSOM employs tomographic imaging, *i.e.,* in contrast to optoacoustic microscopy also out-of-focus signals within the double-conical sensitivity field of the transducer are detected. Therefore, in RSOM image reconstruction has to be employed in order to restore the original 3D optical absorption distribution of the imaged absorbers. Here, we used 3D filtered backprojection where the sensitivity field of the detector was modeled as a hyperboloid with a Gaussian-weighted lateral profile^3^. From the 3D reconstruction, images were created using maximum intensity projections (MIP). For the *Xenopus laevis in vitro* measurement, the thin cell layer was measured at the focal plane of the transducer and no image reconstruction had to be employed. In this case, the photoacoustic signals were cropped around the focal time and directly projected along the depth dimension to form MIPs of the imaged cell layer. Generally, image processing was performed with ImageJ. The same image processing tools were utilized for the control and stimulation data and applied homogeneously across the entire images. The insets were further processed after selection.

### Analysis of the pigmented area with brightfield microscopy and RSOM

The pigmented areas of imaged cells were analyzed using the ROI manager tool from the ‘Fiji’ software^43^. The subsequent statistical analysis was performed in ‘GraphPad Prism 6’ (GraphPad Software, CA, USA). The normality of the data was assessed calculating a Shapiro−Wilk normality test, KS normality test, and D’Agostino & Pearson omnibus test. If data were normal according to at least two of these methods, a *t*-test was used; otherwise, nonparametric tests were employed.

Detailed methods on MORSOM imaging and the frequency analysis can be found in the Supplementary Information.

## ASSOCIATED CONTENT

### Supporting Information

Supplementary Figures S1-S4 and methods (PDF) Supplementary Movies S1 and S2 (AVI)

**Funding Sources** We are grateful for financial support from the European Research Council under grant agreements ERC-St: 311552 (A.L., G.G.W.), the DFG Reinhart Koselleck project “High resolution near-field thermoacoustic sensing and imaging” NT 3/9-1 (D.S., M.O., V.N.) and the Federal Ministry of Education and Research, Photonic Science Germany, Tech2See-13N12624 (D.S., M.O., V.N.).
**ABBREVIATIONS** BF, brightfield; cAMP, cyclic adenosine 3’-5’-monophosphate; CNO, clozapine N-oxide; CPS, cumulative power spectrum; DREADD, designer receptor exclusively activated by designer drugs; FACS, fluorescence-activated cell sorting; FFT, fast Fourier transform; FOV, field of view; GFP, green fluorescent protein; GPCR, G-protein-coupled receptors; MCH, melanin-concentrating hormone; MIP, maximum intensity projection; MORSOM, multi-orientation raster-scan optoacoustic mesoscopy; MSH, melanin-stimulating hormone; NA, numerical aperture; PaPiReS, photoacoustic pigment relocalization sensor; PCR, polymerase chain reaction; PBS, phosphate buffer saline; ROI, region of interest; RSOM, raster-scan optoacoustic mesoscopy;

